# Interface-Sensitive Epi-Coherent Anti-Stokes Raman Scattering Microscopy for Imaging Cell Adhesion Dynamics

**DOI:** 10.1101/2025.10.13.682172

**Authors:** Mingmin Zhou, Bin Dong, Laura Lukov, Seohee Ma, Xinyi He, Chi Zhang

## Abstract

Studying cell adhesion dynamics is critical to understand how cells interact with the extracellular matrix and how they migrate. Conventional microscopy techniques for interfacial studies either lack chemical information or require fluorescence labeling. We found that homodyne epi-coherent anti-Stokes Raman scattering (epi-CARS) microscopy, when focused as interfaces, has intrinsic interfacial selectivity. The interference of the back-reflected nonresonant signals amplifies the contrast of cell adhesion areas, generating a distinctive negative contrast at attachment areas, which reflects the distance between the cell membrane and the substrate with tens of nanometers precision. The Epi-CARS configuration also provides high contrasts for cellular lipid droplets when applied in the inverted microscope configuration. Furthermore, by incorporating a pinhole into our confocal epi-CARS system, we can effectively reject out-of-focus reflections that degrade epi-CARS image quality, allowing us to amplify adhesion and lipid droplet contrasts for complex samples, such as cells. We applied this method to study cell-substrate adhesion dynamics of mitotic cells, revealing adhesion site splitting during mitosis and their outward development during postmitotic spreading. Furthermore, we found that after mitosis, the leading edges exhibited homogeneous adhesion regions with thicker water layers, whereas the retracting edges showed heterogeneous adhesion patterns with thinner layers at the contact sites.

## Introduction

Cell adhesion is an essential process for cells to regulate their activities, such as migration[1], mitosis[2], embryogenesis[3], and cancer metastasis[4]. It is one of the determining factors of how a cell interacts with its neighboring cells and the extracellular matrix (ECM). This adhesion involves a complex interplay of biochemical signaling and mechanical anchoring, mediated by the synthesis and organization of transmembrane adhesion proteins such as integrins[5] and cadherins[6]. These proteins facilitate connections between the cytoskeleton and the ECM, triggering downstream pathways that control motility, survival, and proliferation.

Studying cell adhesion requires imaging techniques that can resolve membrane-substrate interaction with high spatial resolution and interfacial selectivity. Interference reflection microscopy (IRM)[7–9] was the first technique applied for such studies because its contrasts, generated by interference of reflections at the interface, directly reflect the distance between the cell membrane and the substrate. As a label-free, interference-based imaging technique, IRM has no chemical specificity and therefore was largely superseded by fluorescence-based techniques such as total internal reflection fluorescence microscopy (TIRFM)[10–12]. However, TIRFM requires fluorescence labeling, which could cause functional perturbation to biological systems. Furthermore, time-lapse fluorescence imaging is constrained by fluorophores’ photobleaching. In parallel, another label-free approach, typically known as interferometric scattering (iSCAT) microscopy[13–15] today, has extended interference-based imaging techniques with improved sensitivity and axial resolution but remains non-chemical-specific. Furthermore, iSCAT works well on simple low-scattering surfaces, but is challenging to apply in complex biological systems due to interference of background scattering and refractive index variation.

Coherent anti-Stokes Raman scattering (CARS) microscopy enables label-free imaging based on intrinsic vibrational modes of molecules, offering chemical specificity with high acquisition speed[16–19]. However, the nonresonant background, which originates from the electronic response of the medium and propagates mainly in the forward direction, could obscure weak resonant signals and reduce image contrasts. While polarization-sensitive[20], time-resolved [21], and phase-retrieval[22] methods are employed to reduce nonresonant background, these approaches are complicated in the optical setup or data processing.

An alternative strategy is epi-CARS microscopy, which only collects backward CARS signals. This geometry not only minimizes the nonresonant background from the bulk medium[16, 23] but also makes the detection inherently interface-sensitive, as the signal mostly arises from discontinuities in refractive index or nonlinear susceptibility at interfaces[24]. Potma *et al*. discovered the homodyne enhancement of near-interface lipid bilayers through interfering with the back-reflected glass coverslip nonresonant signal and utilized the interference effects of two lipid bilayers to achieve the beyond diffraction-limited resolution of microscopy[25]. Langbein *et al*. demonstrated a dual-polarization epi-heterodyne CARS microscopy that implemented an external reference beam, enabling high-contrast, topographically sensitive imaging of individual lipid bilayers[26]. Furthermore, in an inverted microscopy setup, the epi-CARS geometry positions all excitation and detection optics underneath the sample, enabling direct imaging of cells in culture dishes while accommodating a stage-top incubator. This configuration is particularly advantageous for longitudinal live-cell imaging [27], a task that is typically challenging in coherent Raman microscopy. However, CARS microscopy has not yet been tailored to investigate surface-specific properties for complex systems such as cell adhesion and migration dynamics.

In this study, we present femtosecond epi-CARS microscopy to investigate cell-substrate interactions with high interface selectivity. The homodyne detection scheme provides a much simpler configuration compared to heterodyne detection. In the epi-configuration, we observed a characteristic negative contrast from the cell attachment region, which arises from the difference of the back-reflected interference nonresonant signal at water-substrate and water-membrane interfaces. Such a contrast is very specific to the interface with tens of nanometers selectivity and can be observed in both inverted and upright microscopy configurations. Above the attachment interface, lipid droplets (LDs) within the cells have high contrasts in epi-CARS, a feature not attainable with either IRM or iSCAT. We systematically investigated the contrast of epi-CARS at different depths and using both inverted and upright optical geometries. Furthermore, we found that in the epi-CARS configuration, incorporating a confocal pinhole effectively suppresses out-of-focus reflections and improves image quality. This finding challenges the long-standing assumption that CARS signals originate exclusively from the focal volume and therefore do not require confocal spatial filtering. Finally, we applied this method to monitor cell attachment dynamics during mitosis and observed time-lapse changes in adhesion areas. These include the splitting of adhesion sites during mitosis and the enrichment of adhesion sites at leading edges oriented away from the division center during post-mitotic cell separation. Furthermore, we found that after mitosis, the leading edge exhibits spatially homogeneous adhesion areas between the cell membrane and the substrate, characterized by thicker water layers, whereas the retracting edges show more spatially heterogeneous adhesion site distributions with thinner water layers at the adhered regions. These results demonstrate epi-CARS as an interface-selective and label-free imaging tool for studying adhesion dynamics of live cells with substrates.

## Results

### Contrasts of Femtosecond Epi-CARS for the inverted microscope

The optical system used in this study is illustrated in **Figure 1**, where epi-CARS was implemented in both inverted and upright configurations. This flexibility enables a comprehensive investigation of epi-CARS in both geometries for live-cell imaging. Details of the CARS instrumentation can be found in Experimental Section and an earlier publication.[27] In the inverted configuration, the epi-CARS signals are de-scanned by the same scanning mirrors and can be detected in a confocal setup by introducing a pinhole at the sample-conjugate plane before the detector. In contrast, in the upright configuration, the signals are not de-scanned and are acquired directly by the detector.

**Figure 1.**
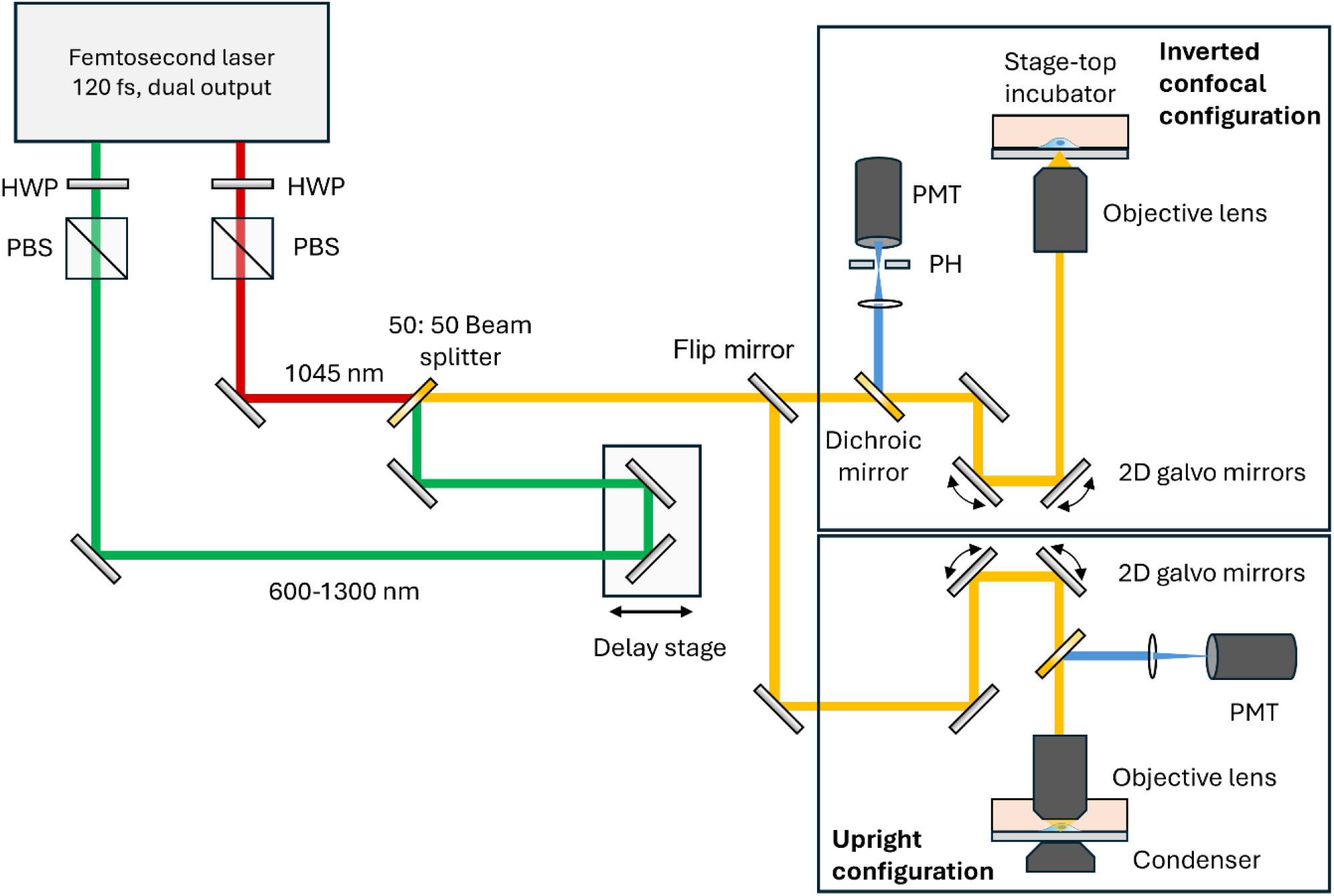
Schematic of femtosecond epi-CARS microscopy used in experiments. The CARS microscopes were implemented in both inverted and upright configurations. In the inverted microscope configuration, the epi-CARS signals are descanned by a 2D galvo scanner before reaching the detector. A 300 µm pinhole can be positioned at the image conjugate plane before the detector to block out-of-plane CARS signal reflections. A stage-top incubator was integrated into the inverted configuration for live cell time-lapse monitoring. HWP, half-wave plate; PBS, polarizing beam splitter cube; PH, pinhole; PMT, photomultiplier.

Here, we first examined epi-CARS in the inverted microscope configuration. Cells were cultured on glass-bottom dishes, where the glass substrate supports cell growth. Unlike polymer substrates, glass does not produce strong resonance signals that could interfere with cellular CARS signals in the C-H stretching region. Glass surfaces are typically weakly negatively charged, allowing proteins from the culture medium to adsorb onto the surface and form an anchoring layer for cells. Cell membrane adhesion molecules, such as integrins, then bind to these adsorbed proteins to mediate attachment. Between the cell membrane and the protein-coated glass surface generally lies an ultrathin water layer.

In homodyne epi-CARS, membrane signals can be detected due to the effective suppression of the nonresonant signals from the medium bulk that mostly travel forward[24]. However, when focused on the cell-substrate interface, a small fraction of the nonresonant signal from the glass substrate can be back-reflected by various interfaces close to the substrate due to linear refractive index discontinuities. Although each reflected portion represents only a small fraction of the forward signal, their amplitudes are usually stronger than those of the backward-generated resonant CARS signals from the membrane lipid bilayer.

When no cell is present, the interface can be approximated as a simple substrate-water configuration. In cell-adhesion regions, the interface generally has a substrate-water-membrane-cytosol configuration, where all layers are located within tens of nanometers above the substrate. Because the refractive index difference between the membrane and cytosol is smaller than that between the membrane and water, the contribution from the membrane-cytosol interface is comparatively weak. Thus, the system can be further simplified to a substrate-water-membrane configuration. In the near-infrared range, the typical refractive indices are approximately 1.51 for the substrate, 1.33 for water, and 1.45 for the membrane at 650 nm CARS signal wavelength.

When only the substrate-water interface is present,

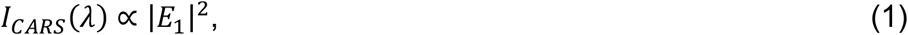

where *I*_*CARS*_ is the epi-CARS intensity and *E*_*1*_ is the reflected electric field by the substrate-water interface.

In cell-adhered regions, the total epi-CARS signal can be estimated as

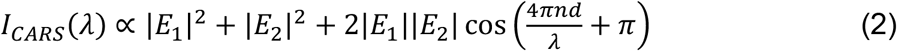

where *E*_*1*_ and *E*_*2*_ are electric fields reflected by the substrate-water and water-membrane interfaces, respectively (**Figure 2A**). The phase π arises from the reflection at the water-membrane interface when entering from the water side (from lower to higher refractive indices). In addition, *n* denotes the refractive index of materials between two interfaces, *d* is the optical path difference, and *λ* is the wavelength.

**Figure 2.**
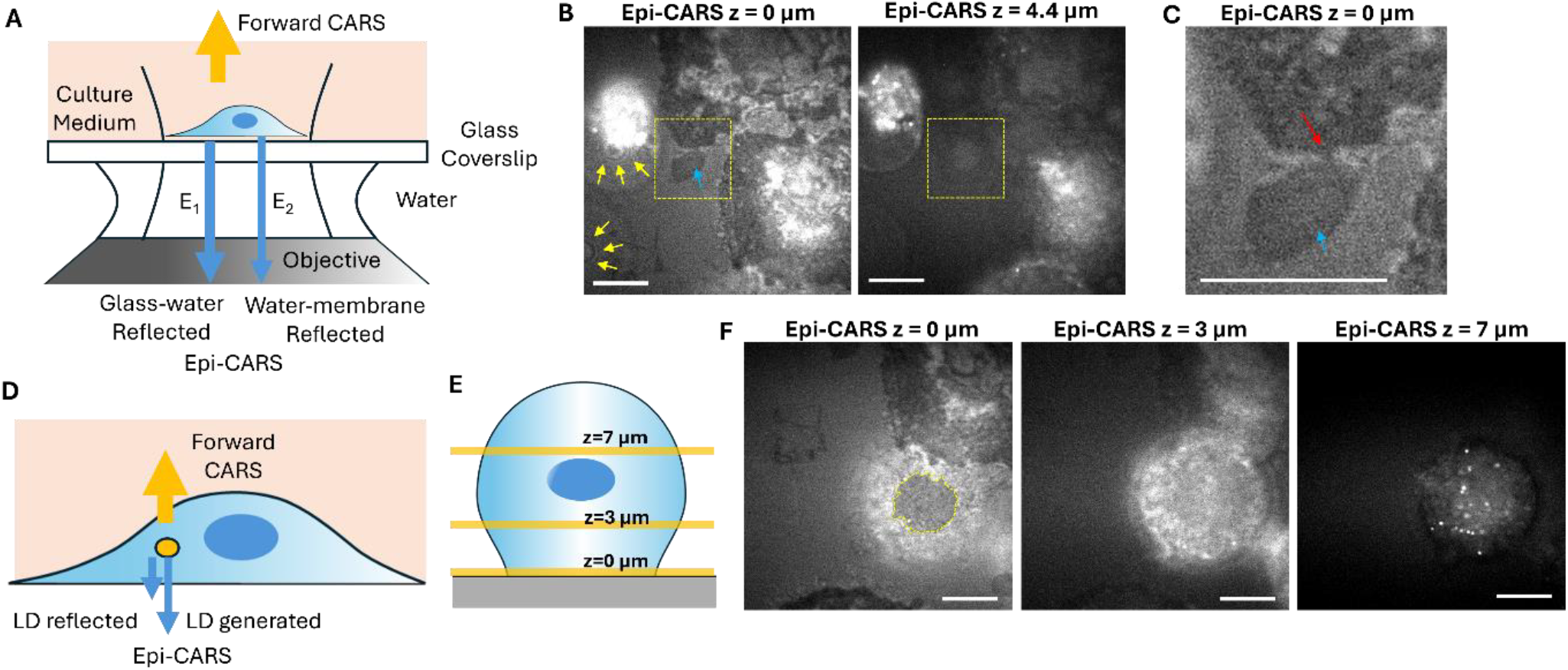
Image contrasts of interface-sensitive femtosecond epi-CARS in the inverted configuration. (A) Schematic of negative contrasts generation in the cell adhesion region via interference of reflected electric fields at the substrate-water and water-membrane interfaces. (B) Epi-CARS images at the cell-substrate interface (z=0 µm) and above the interface (z=4.4 µm) showing various contrast features. The measured vibrational energy is centered at 2899 cm^-1^. The necrotic bleb adhesion sites with the substrate are indicated by the blue arrow. The Newton rings formed by the floating membrane bleb and the substrate are indicated by yellow arrows. (C) Magnified image of the yellow dashed box in panel B, z=0 µm. The necrotic bleb adhesion area and membrane fracture site are indicated by the blue and red arrows, respectively. (D) Schematic of lipid droplet (LD) contrast generation originating from both χ^(1)^ and χ^(3)^ discontinuities. (E) Schematic of imaging a spherical cell at different axial depths (z=0 µm: adhesion plane; z=3 µm: mid-cell; z=7 µm: top cell layer). (F) Epi-CARS images at the three axial depths indicated in panel E. At the cell-substrate interface (z=0 µm), the negative contrast indicates the adhesion area (yellow dashed line). At the top cell layer (z=7 µm), strong LD signals are detected. Scale bars: 10 µm.

To generate a negative contrast at the membrane-glass interface:

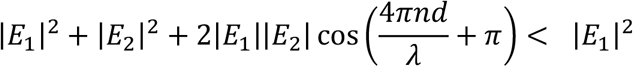

Rearranging this inequality (Supporting information) yields

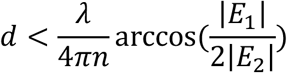

Using typical refractive index values for water, glass, and the membrane (Supporting Information), we can estimate:

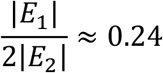

This gives *d* < 51 nm. Thus, when the cell membrane lies within 51 nm of the glass substrate, negative contrast can be generated. Moreover, smaller values of *d* further reduce *I*_*CARS*_, leading to a darker negative contrast.

It is worth noting that in certain non-adhered regions, where the cell membrane is present but sufficiently far from the substrate, the phase 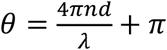 can lead to *I*_*CARS*_ (*λ*) ∝ |*E*_1_|^2^ + |*E*_2_|^2^ + 2|*E*_1_||*E*_2_|*cosθ* > |*E*_1_|^2^, which appears as a positive (brighter) contrast relative to the bare substrate-water interface. A continuous variation in spacing, produced by a spherical surface curving away from the substrate, can generate alternating positive and negative structures similar to Newton rings.

**Figure 2B** shows typical femtosecond epi-CARS images of attached and floating HeLa cells. Here, the confocal pinhole was not implemented, and the cells were fixed using formalin for study. Images were acquired at the substrate-water interface and at higher planes for comparison.

At the interface (z = 0 μm), distinct negative-contrast features indicate regions where the cell is attached to the substrate, particularly along the cell edges. In contrast, the central regions of the cells, where multiple membrane layers are present at higher planes, exhibit brighter signals.

Furthermore, a dead cell with a large necrotic bleb is observed on the left side of the image. This cell is floating and not adhered to the substrate. The large membrane bleb, which contacts the surface, exhibits a characteristic Newton ring pattern around the contact point (yellow arrows). Because the bleb lacks complex membrane layers inside to reflect CARS signals, it displays much weaker signals (blue arrow) overall compared to the host cell. Another membrane bleb is visible at the lower left of the image, showing a similar Newton ring pattern.

The negative contrast of the attached regions disappears rapidly (within 1 μm) as the laser focus is shifted upward. At z = 4.4 μm, the images primarily reveal LDs and ER membranes within the cells. Plasma membrane blebs and floating cell bodies are also more clearly visualized at this axial level. Femtosecond epi-CARS further enables detection of small membrane rupture sites near the interface, as shown in **Figure 2B** (yellow boxed area) and enlarged in **Figure 2C** (red arrow). Such ruptures are likely caused by the formalin fixation process.[28] The adhesion region of the plasma membrane bleb that remains attached to the cell is clearly distinguishable in the z = 0 μm image and further confirmed in the z = 4.4 μm image.

Above the substrate-water interface, the epi-CARS signals primarily arise from CARS generated in the epi-direction. These signals mainly originate from objects smaller than or similar to the focal volume, such as LDs (**Figure 2D)**. When tuned to the CH vibrational region, the epi-CARS response is largely determined by the difference in χ^(3)^ between the object and the surrounding medium |χ^(3)^_obj_ − χ^(3)^_med_|^2^.[24] In contrast, the epi-CARS signal from the pure medium is strongly suppressed.[23] A minor contribution to the signal may also arise from back-reflected forward CARS signals generated by the water or LDs at the bottom or top surfaces of the LDs (**Figure 2D**). This signal is due to χ^(1)^ discontinuity and is less efficient than that from flat surfaces because of LD’s curved geometry.

To demonstrate these contrasts, we imaged a cell in a spherical shape, likely undergoing mitosis, at different axial depths (**Figure 2E**). At the cell-substrate interface, a region of negative contrast indicated cell adhesion, which was much smaller than the overall cell diameter. At z = 3 µm above the interface, the negative contrast disappeared, while the brighter intracellular signals arose from epi-CARS, accompanied by a significant reduction in background. At z = 7 µm, LDs contributed most of the epi-CARS signals, producing very high contrasts. These epi-CARS images demonstrate strong axial sectioning capability, which surpasses that of IRM or iSCAT for complex samples due to the use of tightly focused laser pulses.

It is worth noting that LD contrast varies with the excitation wavelength. According to |χ^(3)^_obj_ − χ^(3)^_med_|^2^, tuning the pump and Stokes lasers to the C-H resonance yields the strongest LD contrast due to the large value of χ^(3)^_LD,R_ and the relatively smaller value of χ^(3)^_water,NR_. When tuned away from both C-H and O-H resonance, the χ^(3)^_LD,NR_ remains greater than χ^(3)^_water,NR._, resulting in weaker but still detectable LD contrast. This contrast is very similar to those detected in third harmonic generation.[29, 30] In contrast, tuning to the water O-H resonance can diminish LD contrast as χ^(3)^_water,R_. approaches or exceeds the value of χ^(3)^ _LD,NR_. Further discussion and figures are provided in Supporting Information and **Figure S1**.

### Effects of Optical Configuration on Epi-CARS Contrast

Similar epi-CARS contrast can be generated at the cell-substrate interface using an upright microscopy geometry. However, this contrast may be masked by strong reflections of forward-CARS signals from the glass substrate bottom. As illustrated in **Figure 3A**, when the dish bottom is exposed directly to air without any medium, a strong reflection arises at the glass-air interface. For thin glass substrates, this reflection can dominate the epi-CARS signals from the cell-substrate interface, resulting in very poor contrasts for adhesion areas (**Figure 3B**), similar to those in forward femtosecond-CARS. This issue can be mitigated by applying immersion oil to the bottom of the dish to match the refractive index of the glass substrate and thereby reduce reflection (**Figure 3C**). Under this configuration, negative adhesion contrasts comparable to those obtained with an inverted microscope can be achieved (**Figure 3D**).

**Figure 3.**
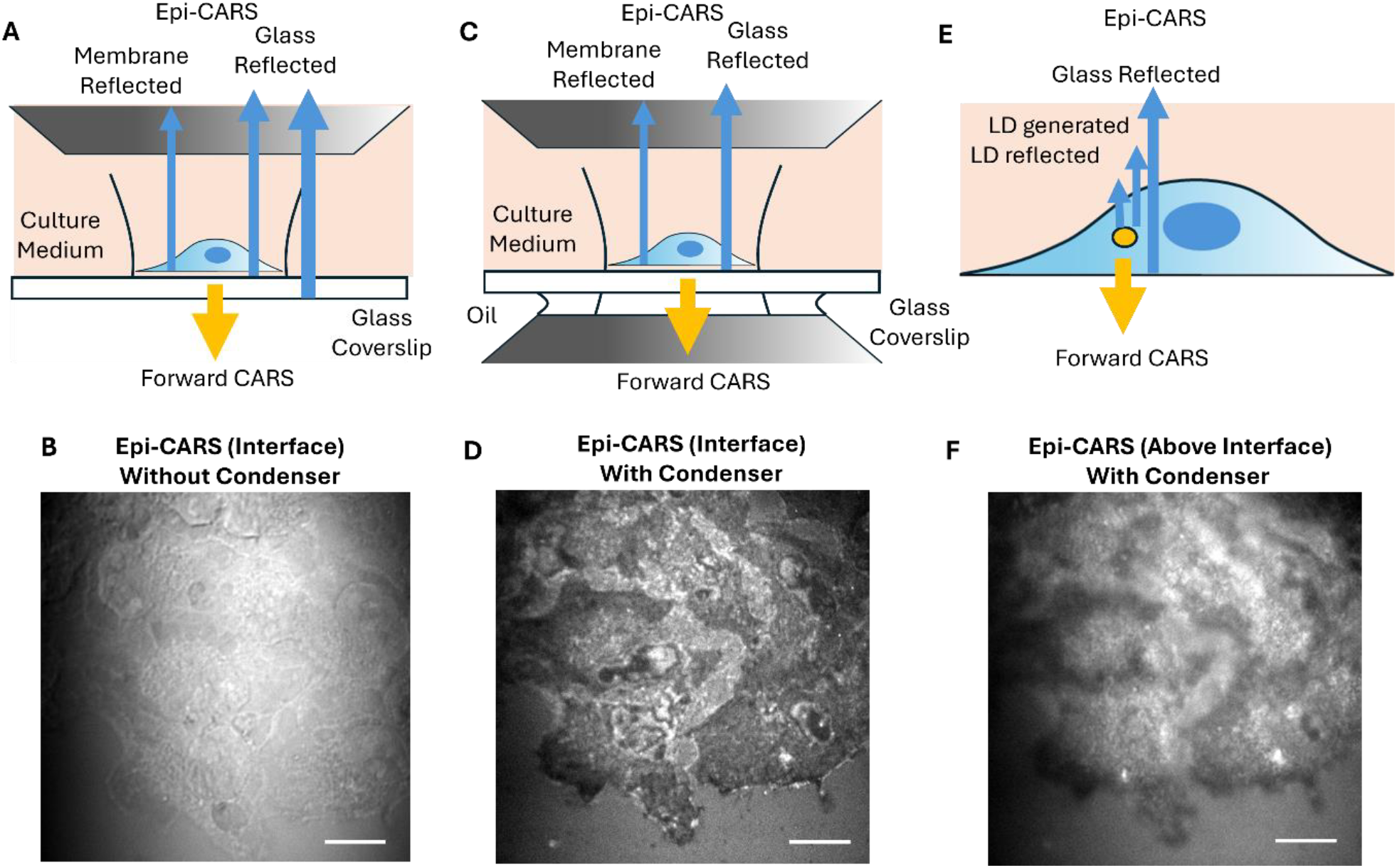
Image contrasts of femtosecond epi-CARS in the upright configuration. (A) Schematic of epi-CARS contrast degradation in cell adhesion areas by strong forward-CARS reflection from substrate-air interface. (B) An epi-CARS image at the cell substrate interface using the geometry in panel A. Signals are dominated by forward CARS reflection. (C) Schematic of interfacial cell adhesion epi-CARS contrast after applying immersion oil to eliminate the glass-air reflection. (D) An epi-CARS image at the cell substrate interface using the geometry in panel C. (E) Schematic of epi-CARS LDs contrasts generation in upright configuration. The strong reflection from substrate-water interface creates a high background that reduces the contrast of LDs. (F) An epi-CARS image above the cell-substrate interface shows poor contrasts of LDs contaminated by the strong forward-CARS reflection. Scale bars: 10 µm.

When the focal plane is shifted into the cell above the substrate interface, LD contrast in the upright configuration is strongly affected by forward nonresonant CARS reflections at the substrate-cell interface (**Figure 3E,F**), showing a blurry image with membrane negative contrasts overlayed. This behavior is different from that in the inverted microscope, where LDs exhibit very high contrasts with minimal background interference.

Furthermore, compared to the upright configuration, the inverted setup keeps the objective lens outside the culture dish and provides a much larger working space for implementing stage-top incubators. Collectively, the inverted geometry is preferred for epi-CARS microscopy.

### Confocal Contrast Enhancement of Epi-CARS

It has long been assumed that CARS signals possess intrinsic axial sectioning capability, since they are generated only at the laser focus, where the energy density is sufficiently high, and thus a confocal pinhole is considered unnecessary.[31] This assumption generally holds for forward signals. However, in the case of epi-CARS, the contrast is significantly affected by the reflections of forward-CARS at different interfaces.[17] As a result, out-of-focus interfacial reflections can still be collected by the large-area photodetector, contaminating the signals from the focal plane. This effect can be effectively suppressed by introducing a confocal pinhole at the conjugate plane of the focal spot, analogous to the principle used in confocal fluorescence microscopy.

In the inverted configuration, the epi-CARS signals were descanned by the galvo scanners. We placed a pinhole before the photomultiplier tube (PMT) to reject contributions from out-of-focus planes. As shown in **Figure 4A**, the addition of the pinhole effectively eliminated reflections from intracellular membranes (from the ER and nucleus) in the outlined regions at the centers of the cells (**Figure 4A**). A more quantitative comparison along selected lines in Figure 4A is shown in **Figure 4B**, showing the rejection of out-of-focus reflections in these areas. As a result, the cell adhesion regions at the cell center retained their negative contrasts, enabling improved characterization of these adhesion areas. In addition, reflections from LDs above the interface were effectively suppressed (red arrows) by the pinhole, leaving only LDs close to the interface visible.

**Figure 4.**
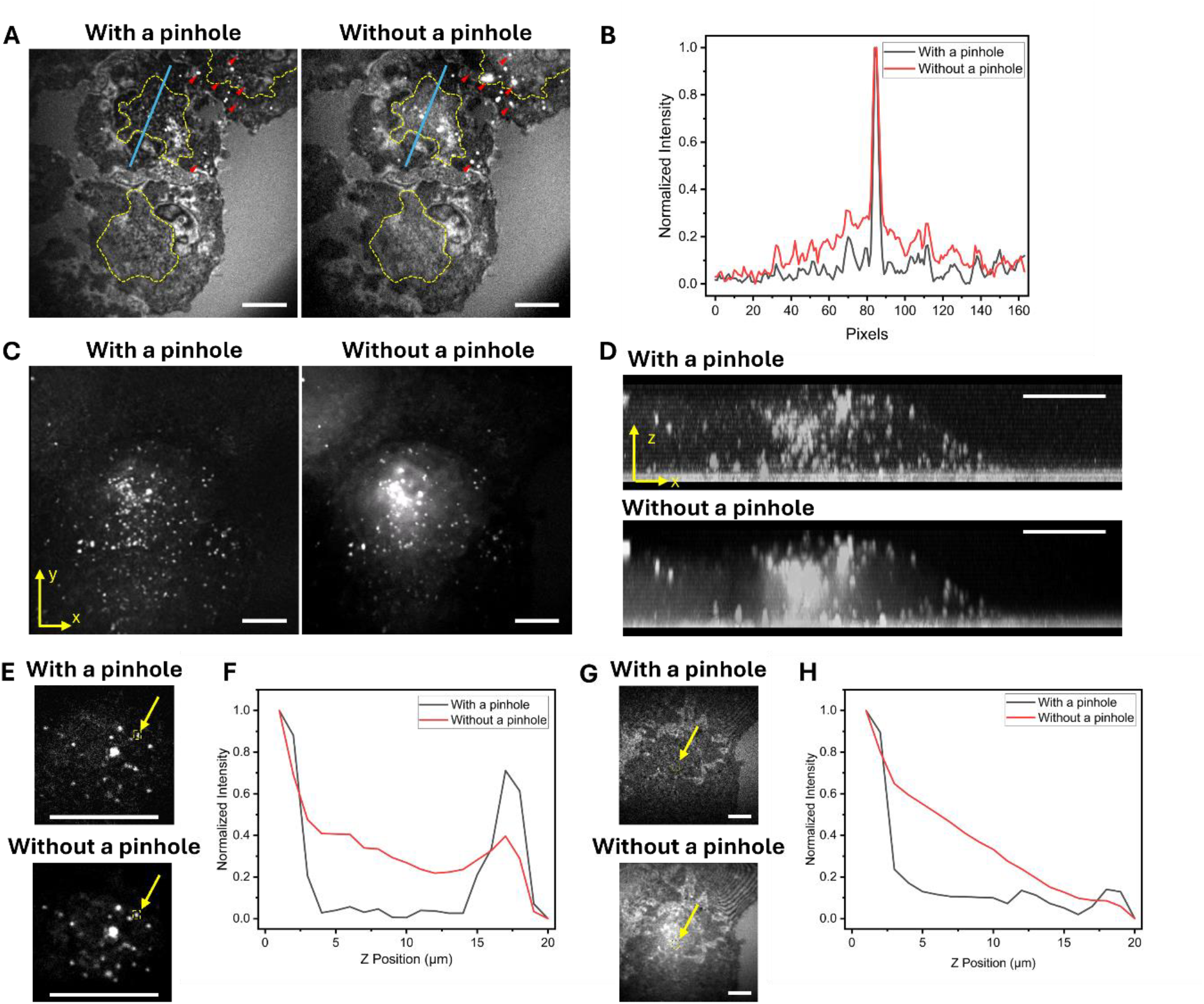
Confocal contrast enhancement of epi-CARS. (A) Epi-CARS images at the cell-substrate interface acquired with (left) and without a confocal pinhole (right). LDs (red triangles) and reflections from the intercellular membrane (yellow dashed outlines) from out-of-focus planes were eliminated when the pinhole was applied. (B) Intensity profiles along blue lines in panel A. (C) Projected epi-CARS images acquired with (left) and without a confocal pinhole (right). Each image is created by summing 15 Epi-CARS images at 1 µm axial interval above the cell-substrate interface. (D) Sagittal (xz) view of the 3D epi-CARS images acquired with (top) and without a confocal pinhole (bottom). (E) Magnified images of a cluster of LDs in the same field of view as panel C, acquired with and without a pinhole. The images are acquired 15 µm above the cell-substrate interface. (F) Epi-CARS axial intensity profiles of an LD (yellow arrow in panel E) acquired with and without a confocal pinhole. (G) Epi-CARS images at the cell-substrate interface acquired with and without a pinhole. (H) Epi-CARS axial intensity profiles of the region of interest (yellow arrow in panel G) acquired with and without a pinhole. Scale bars: 10 µm.

Above the interface, the pinhole effectively suppressed CARS signal reflections from complex intracellular membranes, thus significantly enhancing the contrast of LDs. As shown in **Figure 4C**, without the pinhole, a hazy background appears at the cell center due to reflections from the ER, nuclear, and LDs in other layers, which obscures in in-focus LD contrasts in these regions. With the pinhole applied, individual LDs could be clearly resolved with high contrast and minimal reflection interference. Three-dimensional epi-CARS imaging within this field of view (FOV) was performed, and the sagittal x-z view better illustrates the contrast enhancement of the LDs (**Figure 4D**) by using the confocal pinhole. The corresponding 3D epi-CARS image is provided in Supplementary **Video S1**.

The rejection of surface reflections by the pinhole is best demonstrated by comparing axial intensity profiles. **Figures 4E,F** show the axial profiles of LDs acquired with and without the confocal pinhole. Without the pinhole, the profiles appear asymmetric, with the slope under the LD surface contributed by surface reflections. Introducing the pinhole suppresses these reflections, giving a more symmetric profile with minimal background. Similarly, at the cell-substrate interface, the pinhole effectively rejects forward CARS signal reflections from multiple intracellular membrane layers, thus producing negative contrasts at adhesion regions (**Figure 4G,H**). The axial resolution for LDs is measured to be approximately 1.3-1.4 μm, primarily limited by the axial focal size of the laser (**Figure S2**).

### Investigating Cell Adhesion Dynamics During and After Mitosis

Interface-sensitive epi-CARS imaging is particularly suited for investigating cell adhesion in various conditions. We applied it to examine changes in cell adhesion to glass substrates during mitosis, a fundamental biological process driven by dynamic changes in cell shape and adhesion. Previous studies have shown that adhesion undergoes distinct phases during mitosis, as cells round up between prophase and metaphase, retaining only minimal peripheral adhesions surrounded by filopodia.[2] As cytokinesis proceeds, the centrosome directs cleavage furrow ingression, resulting in daughter cells that subsequently re-establish contact adhesions.[2] These processes can be clearly resolved using interface-sentisitve epi-CARS microscopy. To more precisely identify cells in metaphase, we used HeLa cells transfected with histone-2-mCherry. At this stage, the chromosomes align at the metaphase plate approximately 10 μm above the substrate (**Figure S3**). The cell exhibits a spherical morphology with only a small attachment region at the substrate interface.

Using interface-sensitive epi-CARS imaging, we tracked the sequence of cell adhesion changes during mitosis (selected time points shown in **Figure 5A**). In the early stages, a small region of negative contrast was observed in the cell center, together with small, line-like negative contrasts at the periphery (indicated by yellow triangles), corresponding to filopodia adhesion sites. As cells entered telophase, the central adhesion site split into two major adhesion regions with reduced contact areas, while the peripheral filopodia adhesions remained. These binding sites are highly heterogeneous, accompanied by unattached regions with larger membrane-substrate distances. During cytokinesis, adhesion sites in both daughter cells expanded, reflecting the formation of new integrin-binding areas anchoring the newly divided cells. In the early post-division stage, the daughter cells flattened and re-adhered to the substrate, exhibiting significantly enlarged adhesion regions, particularly extending outward from the division axis (red dashed lines). This oriented adhesion pattern likely facilitates effective separation of the daughter cells and promotes effective reattachment and post-division movement.[19] Such cell adhesion dynamics are illustrated in **Figure 5B**.

**Figure 5.**
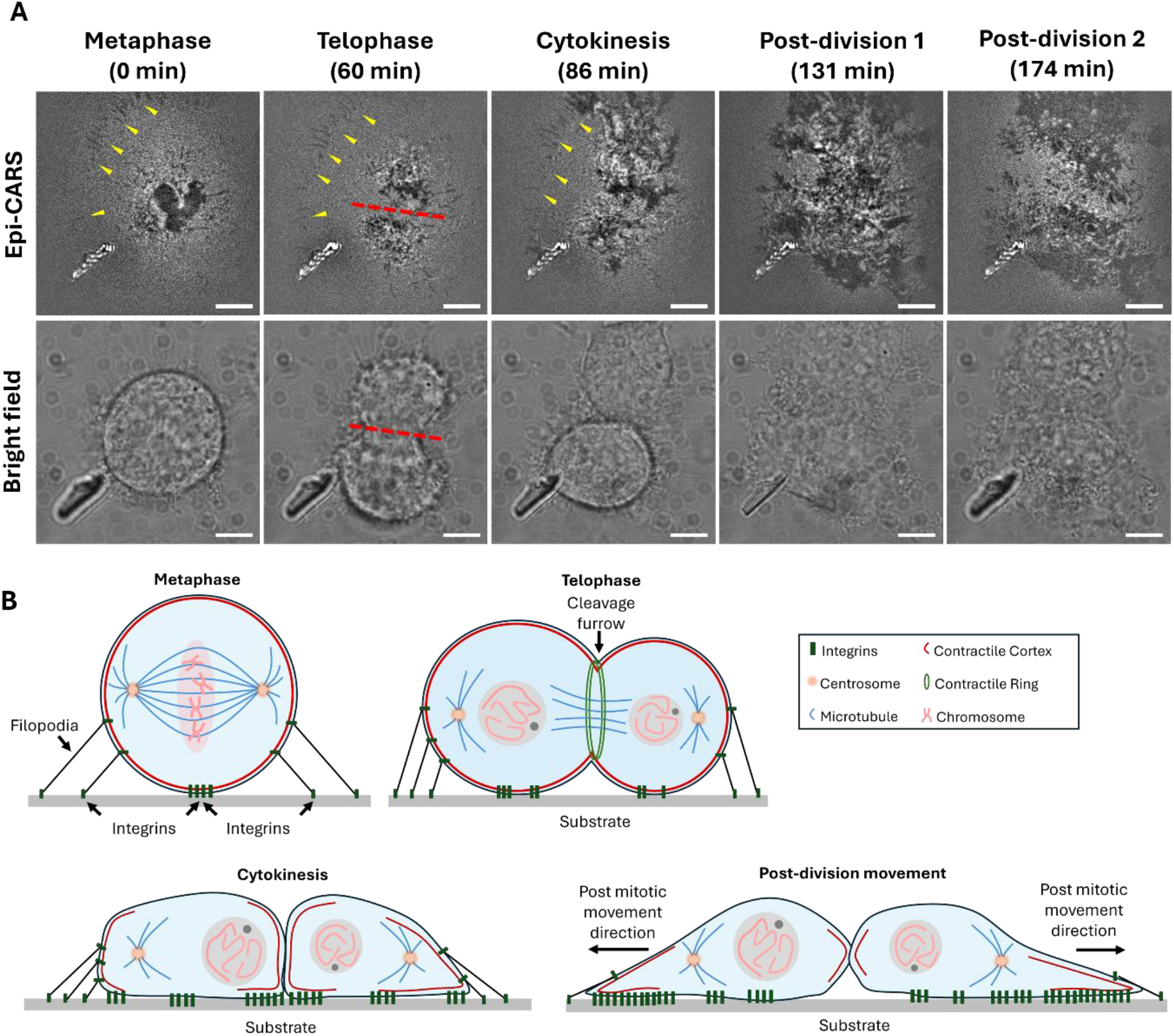
Interface-sensitive epi-CARS imaging reveals cell adhesion dynamics during and after mitosis. (A) Time-lapse epi-CARS images (top) of cell-substrate interface and brightfield images (bottom) from the same field of view for a mitotic cell. Imaging starts in the metaphase. Filopodia adhesion sites are observed from metaphase to cytokinesis and are highlighted by yellow triangles. The division axis is shown in red dashed lines in the telophase. (B) Schematic illustrating the adhesion area changes during mitosis and post-division migration. Scale bars: 5 µm.

Furthermore, by comparing the absolute amplitudes of the negative contrasts at different cell regions, we can infer differences in the distance between the cell membrane and the substrate, indicating the thickness of the intervening water layer. By selecting the epi-CARS signal from the bare glass substrate (background) and from the cell adhesion areas (adhesion), and analyzing the difference in *I*_*background*_ - *I*_*adhesion*_, we quantified and compared the negative contrasts near the division center and at the leading edge during various mitotic stages (**Figure 6A**). The adhering leading edges only began to form after cytokinesis.

**Figure 6.**
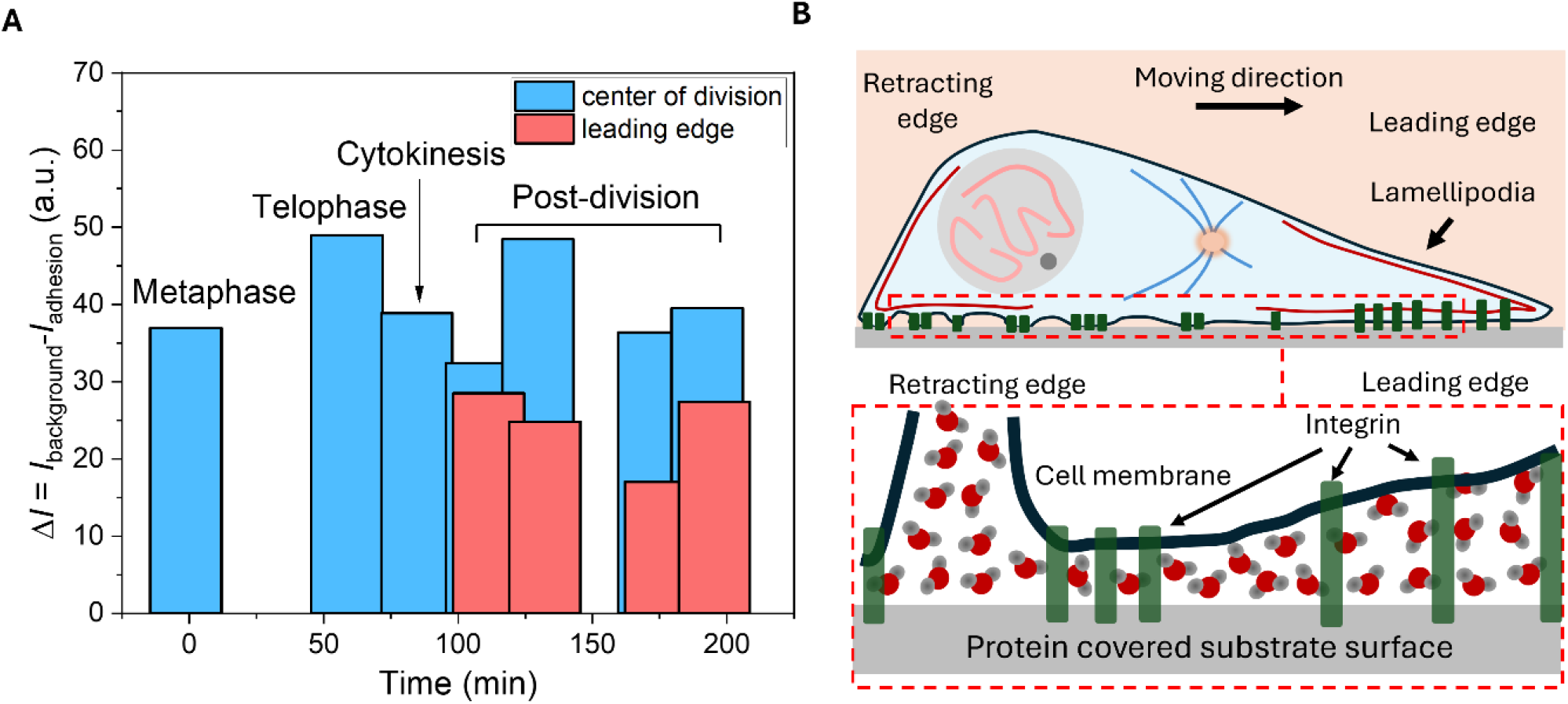
Negative contrasts in the adhesion region reveal variations in membrane-substrate distance and water layer thickness. (A) Analysis of the absolute amplitude of the negative contrast near the division center (blue) and at the leading edges (red) of dividing cells. (B) Schematic illustration showing differences in water layer thickness between the cell membrane and substrate at the retracting and leading edges during post-mitotic movement.

In all cases, the negative contrast amplitude at the division center remained relatively constant and large, while it was significantly smaller at the leading edge. This indicates that the distance between the cell membrane and the substrate at the newly formed adhesion regions in the leading edges is greater than at the contracting edge. The leading edge typically corresponds to the lamellipodium, a protrusive structure that facilitates cell migration. Our results suggest that lamellipodium adhesions form relatively homogeneous adhesion domains but maintain a comparatively thicker water layer in between, still within 51 nm of the substrate. In contrast, at the contracting end, the adhesion sites are more heterogeneous, with regions closely attached to the substrate featuring thinner water layers, interspersed with unadhered regions having membrane-substrate distances exceeding 51 nm (**Figure 6B**).

The observed differences in membrane-substrate distance are likely influenced by variations in integrin conformation.[32–35] The structural states of these integrins could be further investigated using fluorescence resonance energy transfer (FRET) microscopy, which we plan to pursue in future studies. Interface-sensitive epi-CARS has the potential to provide new insights into how interfacial LDs are associated with cell adhesion. Furthermore, our study also demonstrates that interface-sensitive epi-CARS enables real-time monitoring of adhesion dynamics during live-cell division without inducing significant phototoxicity that could disrupt mitosis or trigger apoptosis.

## Conclusion

In summary, we demonstrate femtosecond epi-CARS microscopy as a label-free chemical imaging approach for imaging cell-substrate adhesion and intracellular LDs. At the interface, interference-enhanced contrast provides critical information for measuring the distance between the cell membrane and the glass substrate, enabling quantitative insights into cell adhesion. This interference contrast is highly sensitive to membrane-substrate distance within tens of nanometers above the substrate.

Compared to other interference-based techniques such as IRM and iSCAT, where negative contrasts arise from linear interference and can be extended over relatively large axial distances, the negative contrast in epi-CARS originates from interference of the forward nonlinear optical signal and is highly interface-selective. The implementation of a confocal pinhole effectively suppressed the reflection of out-of-focus interfaces, producing superior optical sectioning, especially for complex samples. While applying a pinhole in traditional IRM produces images with improved qualities, higher-order interference fringes dominate in images taken at higher focal planes [9]. In contrast, interface-sensitive epi-CARS does not produce higher-order fringes. Although CARS microscopy has long been assumed not to require a confocal pinhole due to its intrinsic nonlinear optical sectioning, our results show that in the epi configuration, the pinhole is essential to reject reflections of forward signals from out-of-focus layers and enhance sectioning capability. Furthermore, by comparing inverted and upright geometries, we found that epi-CARS in the inverted configuration is more suitable for studying cell adhesion and intracellular LDs.

Applying epi-CARS, we studied cell adhesion dynamics to the culture dish during mitosis. We found the splitting of the adhesion sites from metaphase to telophase and the enrichment of adhesion areas outward from the division axis to effectively direct post-division movement. Furthermore, our results revealed that during post-mitosis movement, the leading edges of the cells are characterized by spatially homogenous adhesion regions between the cell membrane and the substrate separated by thicker water layers, while the retracting edges show spatially heterogeneous adhesion site distributions and thinner water layers at the adhered regions. These differences are likely driven by variations in actin dynamics and integrin lengths at the leading and retracting edges during cell migration. These findings highlight the capability of epi-CARS to trace live cell adhesion dynamics with minimal phototoxicity.

In this study, we used femtosecond laser pulses, which have high peak power to effectively generate resonant and nonresonant CARS signals. Picosecond lasers could be employed for epi-CARS to generate comparable interfacial-specific contrasts. However, their lower peak power may necessitate longer signal acquisition times or higher pulse energies. The use of pulse-picking techniques could enhance epi-CARS signal generation with picosecond pulses.[36]

Looking ahead, interface-sensitive epi-CARS has strong potential as a platform for studying adhesion-related processes in live cells cultured in 2D. The method is intrinsically compatible with confocal fluorescence microscopy, enabling complementary acquisition of single- or two-photon fluorescence to monitor membrane-specific or intracellular protein dynamics when appropriately labeled. A potential limitation is that the use of femtosecond lasers may cause photobleaching of fluorescent proteins[37], which could restrict long-term monitoring of protein dynamics.

### Experimental Section

#### Femtosecond Epi-CARS Microscopy

The femtosecond epi-CARS microscope configuration is shown in **Figure 1**. To investigate the effect of different microscope geometries on contrast generation, CARS modalities are implemented in both the upright and inverted microscopes. An InSight X3+ femtosecond laser provides synchronized pump (1045 nm) and Stokes (tunable from 690 to 1300 nm) beams at an 80 MHz repetition rate. The laser output provides pulses with a duration of approximately 120 fs. The pulse duration on the sample is about 250 fs. The Stokes laser beam is centered at 1045 nm, while the pump laser beam is tuned to 802 nm to visualize the C-H vibrations. The power at the sample is about 10 mW for both the pump and Stokes beams for live cell imaging. For fixed cell imaging, the pump and Stokes powers were around 15 mW and 20 mW, respectively. Pixel dwell time was set to 10 µs. In both configurations, the beams are combined and focused onto the sample using a 60× water-dipping objective lens (UPlanSApo, 60×/1.2W). A dichroic beam splitter is used to separate and direct the epi-CARS signal to the detector. In the inverted microscope, the epi-CARS signal is de-scanned by the 2D galvo scanner (Saturn-5 system, ScannerMAX) used for laser scanning before being directed into the PMT (H7422-40, Hamamatsu). A confocal pinhole of 300 µm diameter (P300HK, Thorlabs) can be placed at the conjugate plane to reject out-of-focus reflections. A bandpass filter centered at 655 nm (AT655/30, Chroma Technology Corporation) is placed before the PMT to isolate epi-CARS signals. To enable time-lapse monitoring of cell mitosis, a stage-top incubator (WSKMX with STX-CO2O2, Tokai Hit) maintains the physiological condition (37 °C, 5% CO_2_, and 19% O_2_). In the upright microscope, to reduce strong reflection arising from the glass-air interface, an oil condenser and immersion oil can be applied to the dish bottom. Bright-field images of cells are acquired using a CMOS camera (OMAX 10MP).

### Cell Culture

HeLa Kyoto EB3-EGFP cells were maintained in Dulbecco’s Modified Eagle Medium (DMEM, Gibco, Waltham, MA, USA) supplemented with 10% fetal bovine serum (FBS, Gibco, Waltham, MA, USA) and 1% penicillin-streptomycin (Gibco, Waltham, MA, USA). Cells were grown in a humidified incubator at 37 °C with 5% CO_2_. For fixation experiments examining adhesion structures, cells were seeded onto sterilized glass-bottom dishes (MatTek Life Science, Ashland, MA, USA) with 2 mL of culture media. Cells were allowed to adhere overnight and grow to approximately 50% confluency, after which they were either used directly for live-cell imaging or fixed with 10% buffered formalin phosphate (Gibco, Waltham, MA, USA), followed by three phosphate-buffered saline (PBS) washes for subsequent imaging for subsequent epi-CARS imaging.

For live-cell time-lapse monitoring of mitosis, HeLa cells transiently expressing Lamin A-EGFP and histone-2-mCherry were used to visualize chromosomes during cell division. For imaging, cells were maintained at 37 °C with 5% CO_2_ in the stage-top incubator to enable time-lapse cell division monitoring. The two-photon excited fluorescence of mCherry signals was detected by the same epi-CARS channel.

### Data Analysis

All images were processed and analyzed using ImageJ basic functions. For Figure 4B, intensity profiles were extracted along selected lines using the ‘Plot profile’ function. The axial intensity profiles shown in Figure 4F,H from the selected regions of interest (ROIs) were processed using the ‘Plot z-profile’ function and plotted using Origin 2023. The 3D image projected onto the x-y plane was generated by summing 15 consecutive z-stack images acquired at 1 µm axial intervals above the cell-substrate interface. 3D images shown in Video S1 were generated using the ‘3D project’ function in ImageJ. The sagittal (x-z) views were created by rotating the 3D cell image.

For Figure 6, the absolute amplitude of negative contrasts at the division center and leading edge of dividing cells was analyzed by calculating *I*_*background*_ - *I*_*adhesion*_ at the selected regions at different time points. All quantitative data were exported to Origin 2023 for statistical analysis, curve fitting, and figure plotting. The bar graph was plotted using Origin 2023.

## Supporting information

The Supporting Information is available:

Derivation of the Conditions for Negative Contrast in Epi-CARS; Wavelength-Dependent Epi-CARS Contrast; Supporting figures (PDF)

## Author contributions

M.Z. and C.Z. performed the experiment and analyzed the data. B.D. assisted with experiments and optical system maintenance. L.L., S.M., and X.H. contributed to sample preparation and discussion. C.Z. supervised the research and obtained the funding.

## Notes

Dr. Chi Zhang is the founder of Photokinesis LLC, a startup dedicated to commercializing advanced optical control technologies.

## Acknowledge

This research is supported by NIH R35GM147092.

## Data Availability Statement

The data that support the findings of this study are available from thecorresponding author upon reasonable request.

## References

[1] S. SenGupta, C.A. Parent, J.E. Bear, The principles of directed cell migration, Nature Reviews Molecular Cell Biology, 22 (2021) 529–547.

[2] M.C. Jones, J. Zha, M.J. Humphries, Connections between the cell cycle, cell adhesion and the cytoskeleton, Philosophical Transactions of the Royal Society B, 374 (2019) 20180227.

[3] M. Valet, E.D. Siggia, A.H. Brivanlou, Mechanical regulation of early vertebrate embryogenesis, Nature Reviews Molecular Cell Biology, 23 (2022) 169–184.

[4] S.M. Albelda, Role of integrins and other cell adhesion molecules in tumor progression and metastasis, Lab Investigation, 68 (1993) 4–17.

[5] M. Bachmann, S. Kukkurainen, V.P. Hytönen, B. Wehrle-Haller, Cell Adhesion by Integrins, Physiological Reviews, 99 (2019) 1655–1699.

[6] F. Van Roy, G. Berx, The cell-cell adhesion molecule E-cadherin, Cellular and Molecular Life Sciences, 65 (2008) 3756–3788.

[7] A.S.G. Curtis The mechanism of adhesion of cells to glass: a study by interference reflection microscopy, Journal of Cell Biology, 20 (1964) 199–215.

[8] H. Verschueren, Interference reflection microscopy in cell biology: Methodology and applications, Journal of Cell Science, 75 (1985) 279–301.

[9] V.A. Barr, S.C. Bunnell, Interference Reflection Microscopy, Current Protocols in Cell Biology, 45 (2009) 4.23.21–24.23.19.

[10] J.S. Burmeister, L.A. Olivier, W.M. Reichert, G.A. Truskey, Application of total internal reflection fluorescence microscopy to study cell adhesion to biomaterials, Biomaterials, 19 (1998) 307–325.

[11] L. Barbieri, H. Colin-York, K. Korobchevskaya, D. Li, D.L. Wolfson, N. Karedla, F. Schneider, B.S. Ahluwalia, T. Seternes, R.A. Dalmo, M.L. Dustin, D. Li, M. Fritzsche, Two-dimensional TIRF-SIM–traction force microscopy (2D TIRF-SIM-TFM), Nature Communications, 12 (2021) 2169.

[12] K. Affannoukoué, S. Labouesse, G. Maire, L. Gallais, J. Savatier, M. Allain, M. Rasedujjaman, L. Legoff, J. Idier, R. Poincloux, F. Pelletier, C. Leterrier, T. Mangeat, A. Sentenac, Super-resolved total internal reflection fluorescence microscopy using random illuminations, Optica, 10 (2023) 1009–1017.

[13] J.-S. Park, I.-B. Lee, H.-M. Moon, J.-H. Joo, K.-H. Kim, S.-C. Hong, M. Cho, Label-free and live cell imaging by interferometric scattering microscopy, Chemical Science, 9 (2018) 2690–2697.

[14] J.-S. Park, I.-B. Lee, H.-M. Moon, J.-S. Ryu, S.-Y. Kong, S.-C. Hong, M. Cho, Fluorescence-Combined Interferometric Scattering Imaging Reveals Nanoscale Dynamic Events of Single Nascent Adhesions in Living Cells, The Journal of Physical Chemistry Letters, 11 (2020) 10233–10241.

[15] J.-S. Park, I.-B. Lee, H.-M. Moon, S.-C. Hong, M. Cho, Long-term cargo tracking reveals intricate trafficking through active cytoskeletal networks in the crowded cellular environment, Nature Communications, 14 (2023) 7160.

[16] J.-X. Cheng, A. Volkmer, X.S. Xie, Theoretical and experimental characterization of coherent anti-Stokes Raman scattering microscopy, Journal of the Optical Society of America B, 19 (2002) 1363–1375.

[17] C.L. Evans, E.O. Potma, M. Puoris’haag, D. Côté, C.P. Lin, X.S. Xie, Chemical imaging of tissue in vivo with video-rate coherent anti-Stokes Raman scattering microscopy, Proceedings of the National Academy of Sciences of the United States of America, 102 (2005) 16807–16812.

[18] J.L. Suhalim, J.C. Boik, B.J. Tromberg, E.O. Potma, The need for speed, Journal of Biophotonics, 5 (2012) 387–395.

[19] C.L. Dix, H.K. Matthews, M. Uroz, S. McLaren, L. Wolf, N. Heatley, Z. Win, P. Almada, R. Henriques, M. Boutros, The role of mitotic cell-substrate adhesion re-modeling in animal cell division, Developmental Cell, 45 (2018) 132–145.e133.

[20] J.-X. Cheng, L.D. Book, X.S. Xie, Polarization coherent anti-Stokes Raman scattering microscopy, Optics Letters, 26 (2001) 1341–1343.

[21] A. Volkmer, L.D. Book, X.S. Xie, Time-resolved coherent anti-Stokes Raman scattering microscopy: Imaging based on Raman free induction decay, Applied Physics Letters, 80 (2002) 1505–1507.

[22] Y. Liu, Y.J. Lee, M.T. Cicerone, Broadband CARS spectral phase retrieval using a time-domain Kramers-Kronig transform, Optics Letters, 34 (2009) 1363–1365.

[23] A. Volkmer, J.-X. Cheng, X. Sunney Xie, Vibrational Imaging with High Sensitivity via Epidetected Coherent Anti-Stokes Raman Scattering Microscopy, Physical Review Letters, 87 (2001) 023901.

[24] E. Potma, Foundations of Nonlinear Optical Microscopy, Wiley 2024.

[25] E.O. Potma, X.S. Xie, Detection of single lipid bilayers with coherent anti-Stokes Raman scattering (CARS) microscopy, Journal of Raman Spectroscopy, 34 (2003) 642–650.

[26] W. Langbein, D. Regan, I. Pope, P. Borri, Invited Article: Heterodyne dual-polarization epi-detected CARS microscopy for chemical and topographic imaging of interfaces, APL Photonics, 3 (2018) 092402.

[27] G.A. Gonzalez, E.U. Osuji, N.C. Fiur, M.G. Clark, S. Ma, L.L. Lukov, C. Zhang, Alteration of Lipid Metabolism in Hypoxic Cancer Cells, Chemical & Biomedical Imaging, 3 (2025) 25–34.

[28] S. Zhao, H. Liao, M. Ao, L. Wu, X. Zhang, Y. Chen, Fixation-induced cell blebbing on spread cells inversely correlates with phosphatidylinositol 4,5-bisphosphate level in the plasma membrane, FEBS Open Bio, 4 (2014) 190–199.

[29] G.J. Tserevelakis, E.V. Megalou, G. Filippidis, B. Petanidou, C. Fotakis, N. Tavernarakis, Label-free imaging of lipid depositions in C. elegans using third-harmonic generation microscopy, PloS One, 9 (2014) e84431.

[30] T. Pajić, N.V. Todorović, M. Živić, S.N. Nikolić, M.D. Rabasović, A.H. Clayton, A.J. Krmpot, Label-free third harmonic generation imaging and quantification of lipid droplets in live filamentous fungi, Scientific Reports, 12 (2022) 18760.

[31] W. Min, C.W. Freudiger, S. Lu, X.S. Xie, Coherent Nonlinear Optical Imaging: Beyond Fluorescence Microscopy, Annual Review of Physical Chemistry, 62 (2011) 507–530.

[32] M.H. Jo, J. Li, V. Jaumouillé, Y. Hao, J. Coppola, J. Yan, C.M. Waterman, T.A. Springer, T. Ha, Single-molecule characterization of subtype-specific β1 integrin mechanics, Nature Communications, 13 (2022) 7471.

[33] T. Schürpf, T.A. Springer, Regulation of integrin affinity on cell surfaces, The EMBO Journal, 30 (2011) 4712–4727.

[34] P. Stanley, A. Smith, A. McDowall, A. Nicol, D. Zicha, N. Hogg, Intermediate_water_affinity LFA_water_ 1 binds α_water_actinin_water_1 to control migration at the leading edge of the T cell, The EMBO Journal, 27 (2008) 62–75.

[35] J. Zhu, B.-H. Luo, T. Xiao, C. Zhang, N. Nishida, T.A. Springer, Structure of a complete integrin ectodomain in a physiologic resting state and activation and deactivation by applied forces, Molecular Cell, 32 (2008) 849–861.

[36] M.G. Clark, G.A. Gonzalez, C. Zhang, Pulse-picking multimodal nonlinear optical microscopy, Analytical Chemistry, 94 (2022) 15405–15414.

[37] G. Wang, L. Li, J.E. Sorrells, J. Chen, H. Tu, Gentle Label-Free Nonlinear Optical Imaging Relaxes Linear-Absorption-Mediated Triplet, Advanced Science, 12 (2025) e15648.

