## Supporting Information for "Interface-Sensitive Epi-Coherent Anti-Stokes Raman Scattering Microscopy for Imaging Cell Adhesion Dynamics"

### 1. Derivation of the Conditions for Negative Contrast in Epi-CARS

The fraction of back-reflected forward electric fields at the glass-water and water-membrane interfaces can be estimated through the Fresnel equation (0-degree incidence angle):

$$R = \left(\frac{n_1 - n_2}{n_1 + n_2}\right)^2, \qquad T = \frac{4n_1 n_2}{(n_1 + n_2)^2}$$

where $n_1$ and $n_2$ are the refractive indices of entering and existing media. In our experiment, with a pump beam at 802 nm and a Stokes beam at 1045 nm, the CARS wavelength is calculated as 650 nm. Using $n_{air} = 1.00$, $n_{glass} = 1.51$, $n_{water} = 1.33$, and $n_{membrane} = 1.45$, at this wavelength, we obtain:

1) Reflectance at glass-air interface is 4.13%.
2) Reflectance at glass-water interface is 0.40%. Transmission at glass-water interface is 99.6%.
3) Reflectance at water-membrane interface is 0.19%.

The much larger reflectance at glass-air interface accounts for the poor contrast of cell adhesion regions in the upright geometry when the substrate bottom is directly exposed in air. In contrast, the comparable reflectance at glass-water and water-membrane interfaces enables observable interference to modulate the epi-CARS signal at the cell-substrate interface.

When only the substrate-water interface is present,

$$I_{CARS}(\lambda) \propto |E_1|^2$$

where $I_{CARS}$ is the epi-CARS intensity and $E_1$ is the reflected electric field by the substrate-water interface. We define $E_2$ as the reflected electric fields reflected by the water-membrane interface. Using the estimated reflectance and transmittance of two interfaces, we can calculate:

$$\frac{|E_1|}{2|E_2|} \approx 0.24$$

In cell-adhered regions, the total epi-CARS signal can be estimated as:

$$I_{CARS}(\lambda) \propto |E_1|^2 + |E_2|^2 + 2|E_1||E_2|\cos\left(\frac{4\pi nd}{\lambda} + \pi\right)$$

Here, $n$ denotes refractive index of materials between two interfaces, $d$ is the optical path difference, $\lambda$ is the wavelength. A negative contrast is defined as a total intensity lower than the intensity from the glass-water interface alone ($I < |E_1|^2$). This gives

$$|E_1|^2 + |E_2|^2 + 2|E_1||E_2|\cos\left(\frac{4\pi nd}{\lambda} + \pi\right) < |E_1|^2$$

$$|E_2|^2 + 2|E_1||E_2|\cos\left(\frac{4\pi nd}{\lambda} + \pi\right) < 0$$

$$|E_2|^2 < -2|E_1||E_2|\cos\left(\frac{4\pi nd}{\lambda} + \pi\right)$$

Using $\cos(\theta + \pi) = -\cos(\theta)$, equation simplifies into:

$$|E_2|^2 < 2|E_1||E_2|\cos\left(\frac{4\pi nd}{\lambda}\right)$$

Since $|E_1|, |E_2| > 0$, we can divide both sides by $2|E_1||E_2|$ :

$$\frac{|E_1|}{2|E_2|} < \cos\left(\frac{4\pi nd}{\lambda}\right)$$

Therefore, for negative contrasts to occur, the distance between the membrane and the glass must satisfy:

$$d < \frac{\lambda}{4\pi n}\arccos\left(\frac{|E_1|}{2|E_2|}\right)$$

Using the ratio of $\frac{|E_1|}{2|E_2|} \approx 0.24$ and $n_{water} = 1.33$, the upper boundary for the cell-substrate distance that gives a negative contrast is 51 nm. A smaller value in *d* would lead to a lower signal level and darker negative contrast.

### 2. Wavelength-Dependent Epi-CARS Contrast

For a small objects like lipid droplets (LDs), epi-CARS signal is generated at big $\chi^{(3)}$ discontinuities between the object and the surrounding medium. The detected signal intensity is proportional to:[S1]

$$I_{CARS} \propto \left|\chi^{(3)}\text{obj} - \chi^{(3)}\text{med}\right|^2$$

Where $\chi^{(3)}$ has both the resonant and nonresonant distributions: $\chi^{(3)} = \chi_R{}^{(3)} + \chi_{NR}{}^{(3)}$. The epi-CARS contrast is highly wavelength dependent due to the spectral profile of these $\chi^{(3)}$ component (**Figure S1**).

1) On C-H resonance (2899 cm$^{-1}$): $\chi^{(3)}\text{obj} \approx \chi_{LD,R}{}^{(3)}$ is maximized, while $\chi^{(3)}\text{med} = \chi_{water,NR}{}^{(3)}$ is low, leading to a large $\left|\chi^{(3)}\text{obj} - \chi^{(3)}\text{med}\right|^2$. This yields strong positive LD contrasts.
2) Off C-H and O-H resonance (2625 cm$^{-1}$): Here $\chi^{(3)}\text{obj} = \chi_{LD,NR}{}^{(3)}$, $\chi^{(3)}\text{med} = \chi_{water,NR}{}^{(3)}$. Since $\chi_{LD,NR}{}^{(3)} > \chi_{water,NR}{}^{(3)}$, LD contrasts are reduced but still visible. This is similar to the third-harmonic generation (THG) imaging condition.
3) On O-H resonance (3333 cm$^{-1}$): Here $\chi^{(3)}\text{med} \approx \chi_{water,R}{}^{(3)}$, which can be on the similar level as $\chi_{LD,NR}{}^{(3)}$. The $\chi^{(3)}$ discontinuity is small, and therefore the LD contrast is diminished.

In comparison, at the cell-substrate interface, the negative contrast has less impact since it is mostly contributed by the $\chi^{(1)}$ discontinuity at the interfaces.

### 3. Supporting Figures

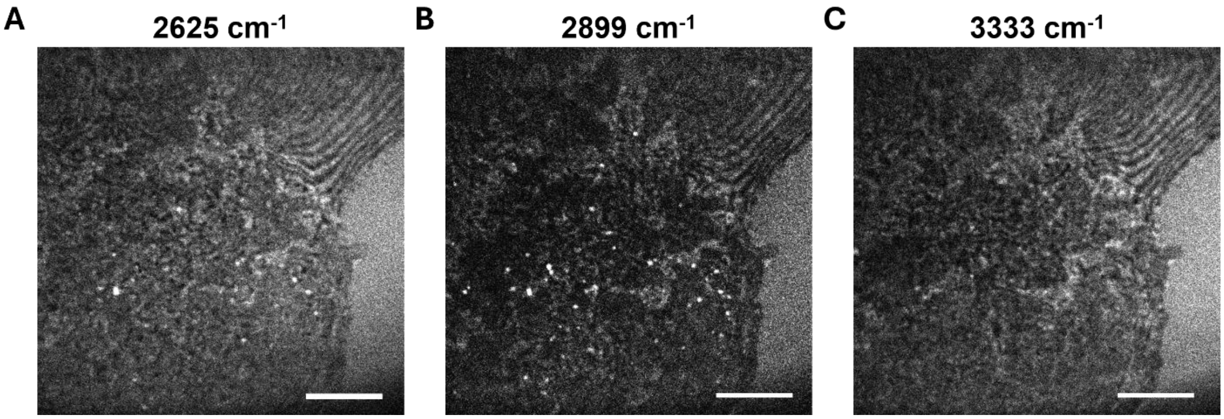


**Figure S1.** Epi-CARS images of cell-substrate interface acquired at different Raman shifts. Images were taken at (A) 2625 $cm^{-1}$ (non-resonant region), (B) 2899 $cm^{-1}$ (C-H stretching resonance), and (C) 3333 $cm^{-1}$ (O-H stretching resonance). Scale bars: 10 μm.

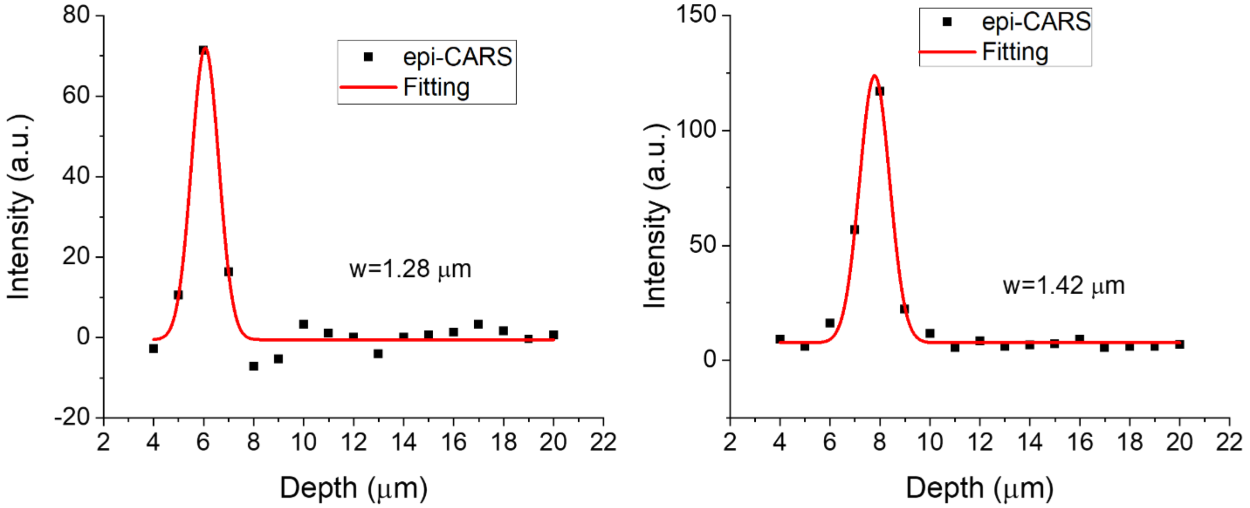


**Figure S2.** Epi-CARS axial intensity profiles of two small lipid droplets selected from the cells in Figure 4C. The profiles were fitted using Gaussian functions. The axial resolution of LDs is estimated to be around 1.3-1.4 μm.

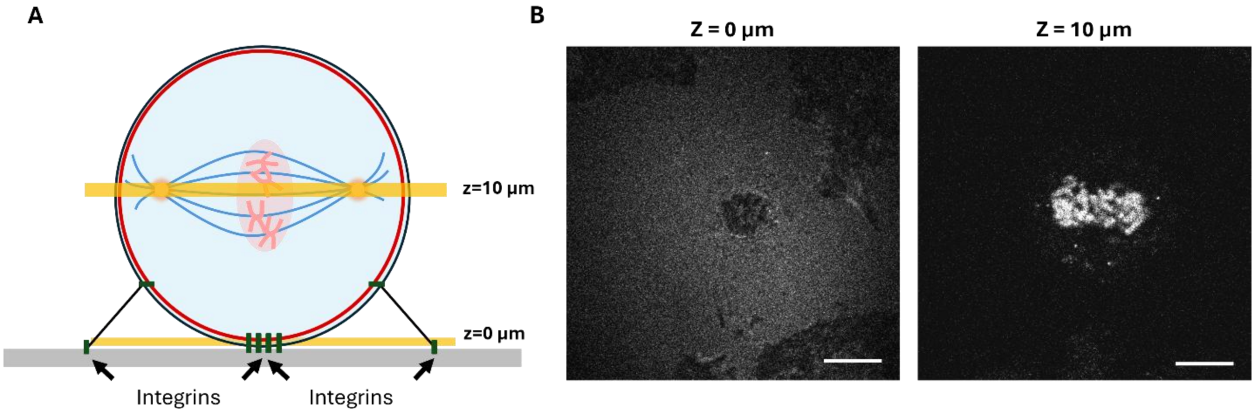


**Figure S3.** CARS and fluorescence imaging of a mitotic cell in the metaphase. (A) An illustration of imaging a mitotic cell at different axial layers in the metaphase. (B) An epi-CARS image (left) of a dividing cell during metaphase at the cell-substrate interface (0 μm) showing the adhesion region. The corresponding two-photon excitation fluorescence image (right) of histone-2-mCherry signals at the layer 10 μm above the interface showing chromosomes structures in the metaphase. Scale bars: 10 μm.

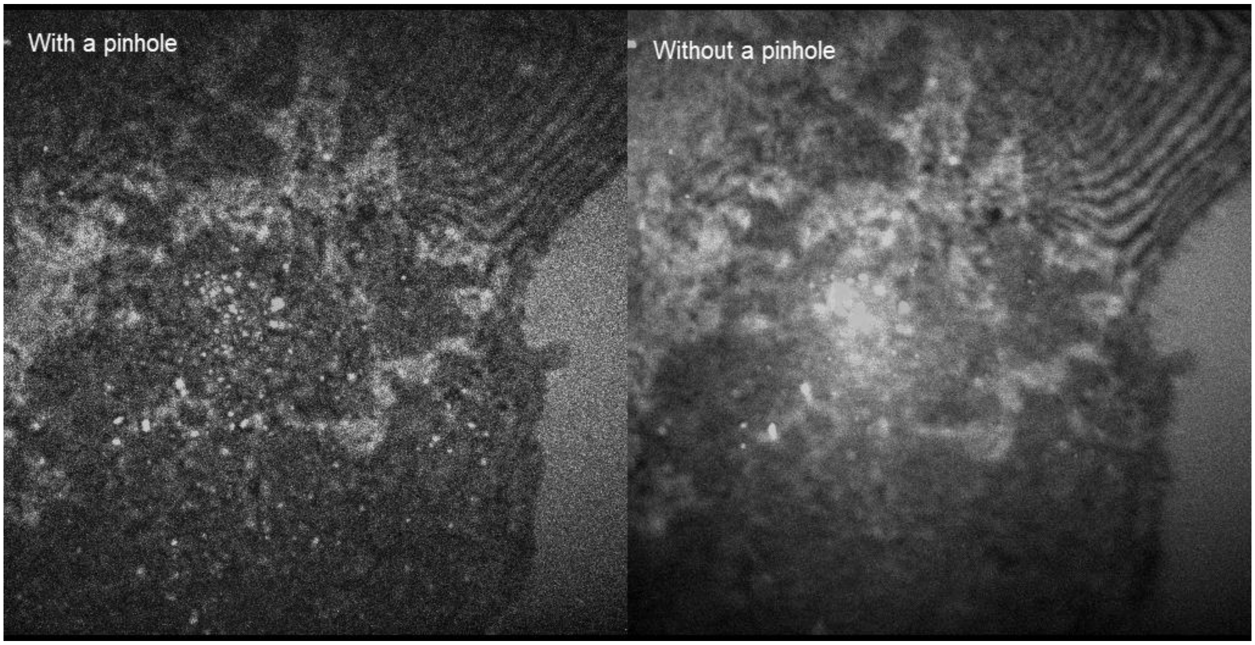


**Video S1.** Comparing 3D epi-CARS images with and without a confocal pinhole.

### References

S1. Potma, E. Foundations of Nonlinear Optical Microscopy; Wiley, 2024.
